## Supplementary material for "Assembly of Synaptic Protein-DNA Complexes: Critical Role of Non-Specific Interactions": complete_SI

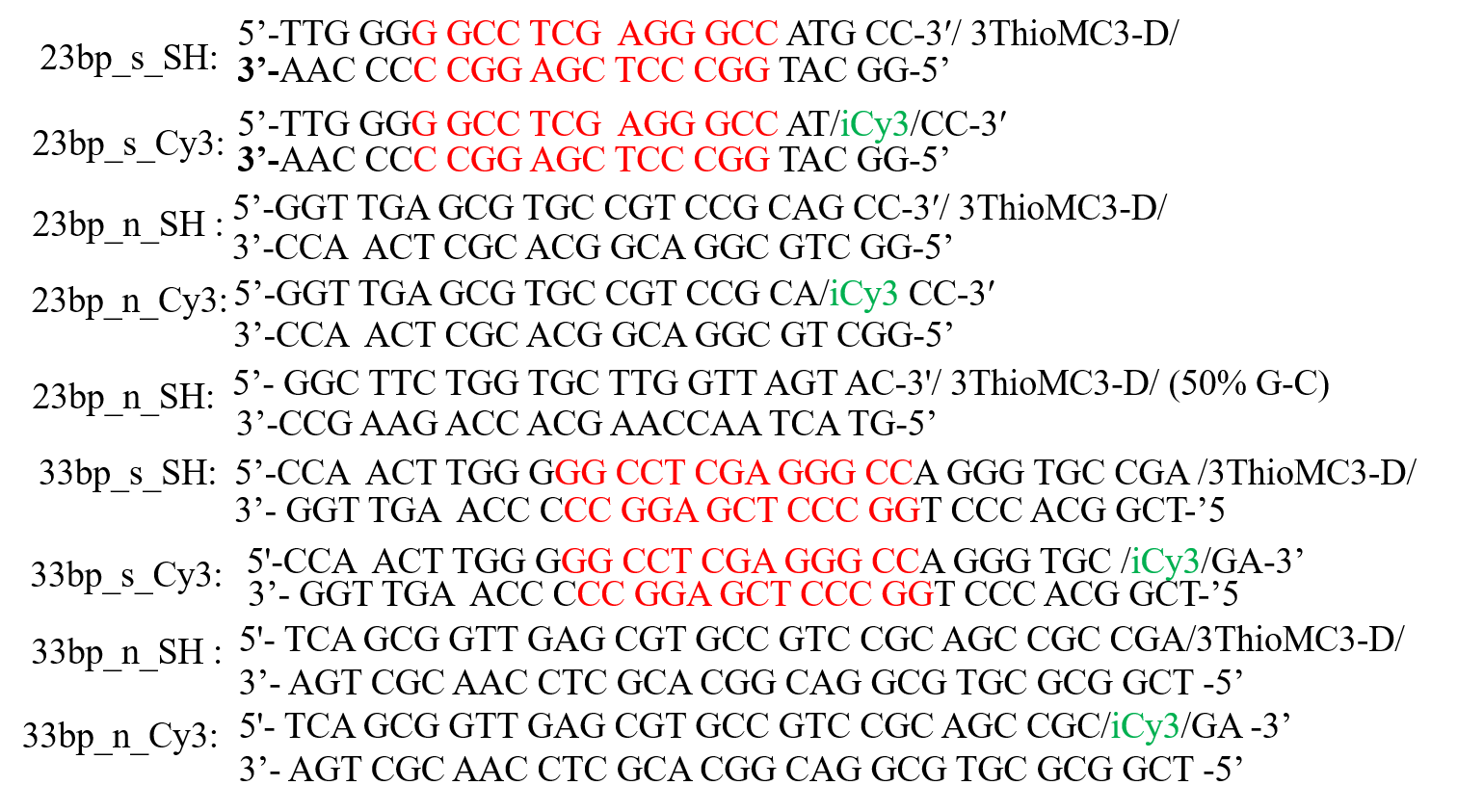

**Figure S1.** Sequence of the 23 and 33 bp DNA substrates used in the study. SfiI cognate sites have been highlighted in red. The letter ‘s’ in the substrate names denotes sequences with the cognate site for SfiI. The letter ‘n’ denotes sequences with no specific site for SfiI. The thiol modification was 3ThioMCD3-D and the internal Cy3 label is indicated as iCy3 in green.

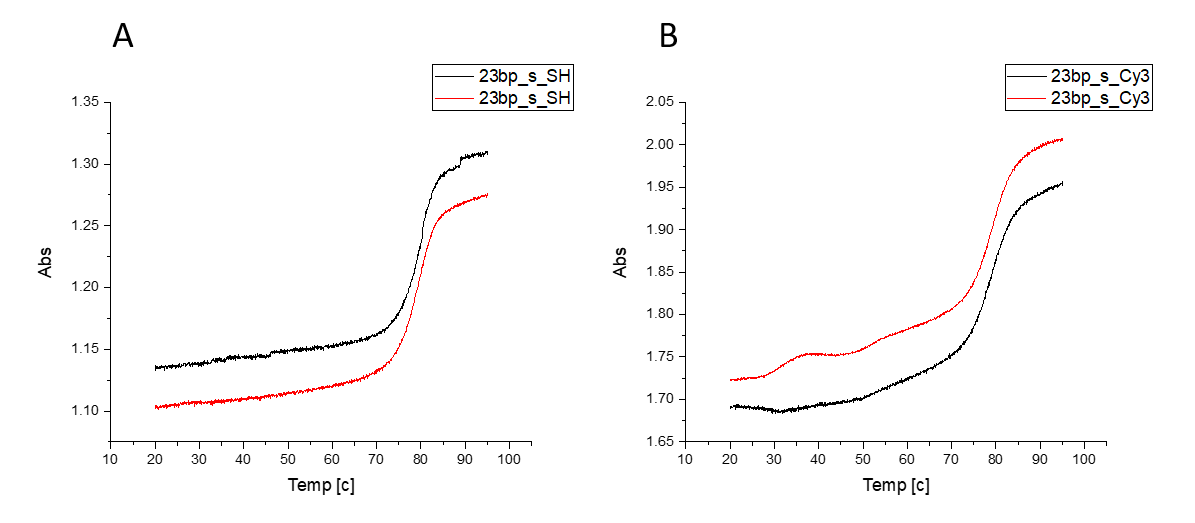

**Figure S2.** UV melting curves for the specific 23 bp DNA duplexes with (A), thiol (SH) and (B), Cy3 modifications. Each melting experiment was performed in duplicates and proper annealing was confirmed from the graphs.

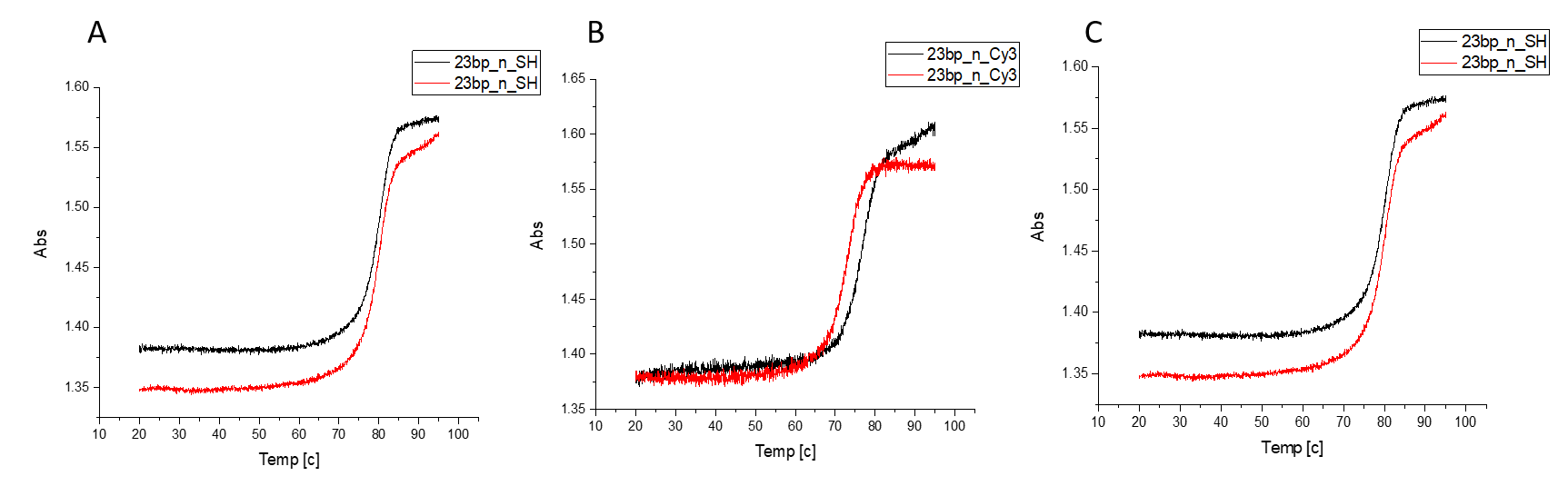

**Figure S3**. UV melting curves non-specific 23 bp DNA duplexes with (A), thiol (SH), (B) Cy3 modifications. (C) Melting curve of DNA duplex with 50% GC content. All the melting curves were performed in duplicates and proper annealing was confirmed from the graphs.

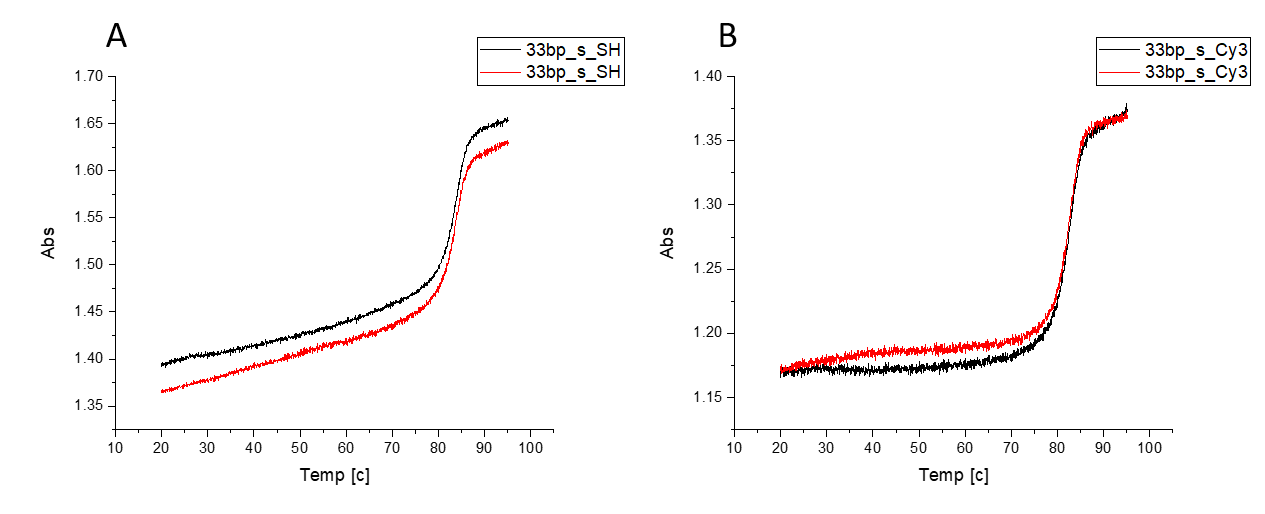

**Figure S4.** UV melting curves for 33 bp specific DNA duplexes with (A), thiol (SH) and (B), Cy3 modifications. All the melting curves were performed in duplicates and proper annealing was confirmed from the graphs.

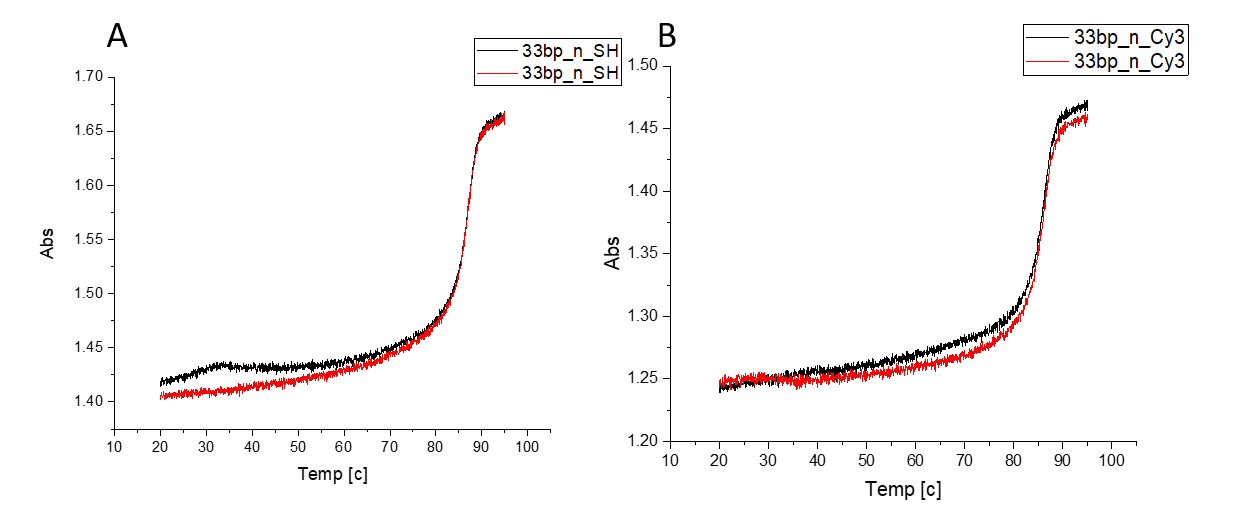

**Figure S5.** UV melting curves of the of the 33 bp non-specific DNA duplexes with (A), thiol (SH) and (B), Cy3 modifications. All the melting curves were performed in duplicates and proper annealing was confirmed from the graphs.

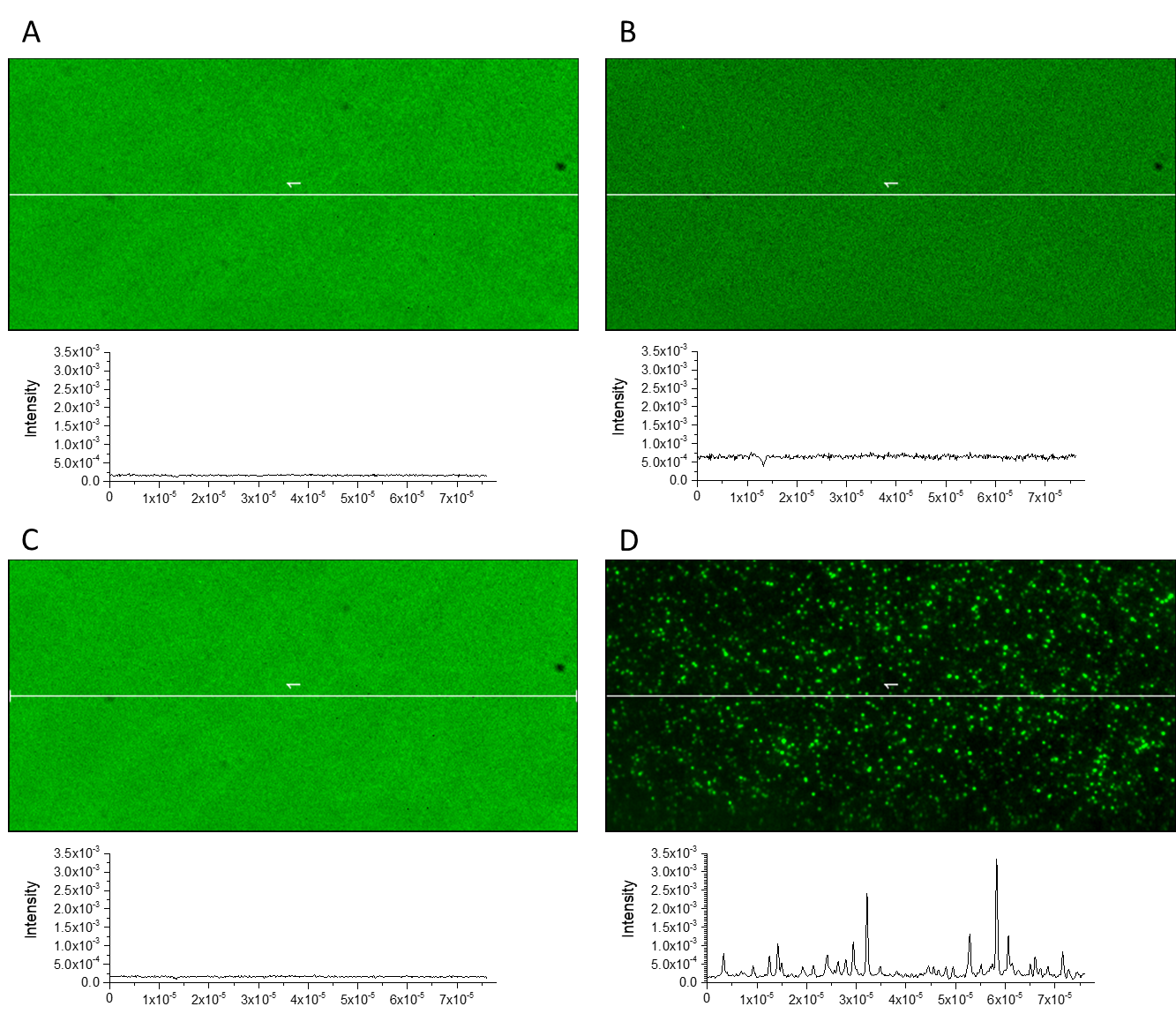

**Figure S6.** TIRF images of surfaces used during TAPIN experiments. (A) Representative image of surfaces with the tethered DNA that was bleached for 30 mins before image acquisition. The roughness (RMS-Sq) of surface is 15.07. (B) Represents the surface in A, after addition of 1 nM SfiI. The roughness (RMS-Sq) of surface is 43.379. (C) Represents a surface like in A after the addition of Cy3-labelled DNA. The roughness (RMS-Sq) of surface is 13.87. (D) Represents a surface like in A with 1nM SfiI and 1 nM of Cy3-labelled DNA, and shows formation of complexes. The roughness (RMS-Sq) of surface is 243.2. All the images (A-D) are the average of the 2000 frames captured at 100ms/frame. The intensity profiles for each scenario is displayed below each image.

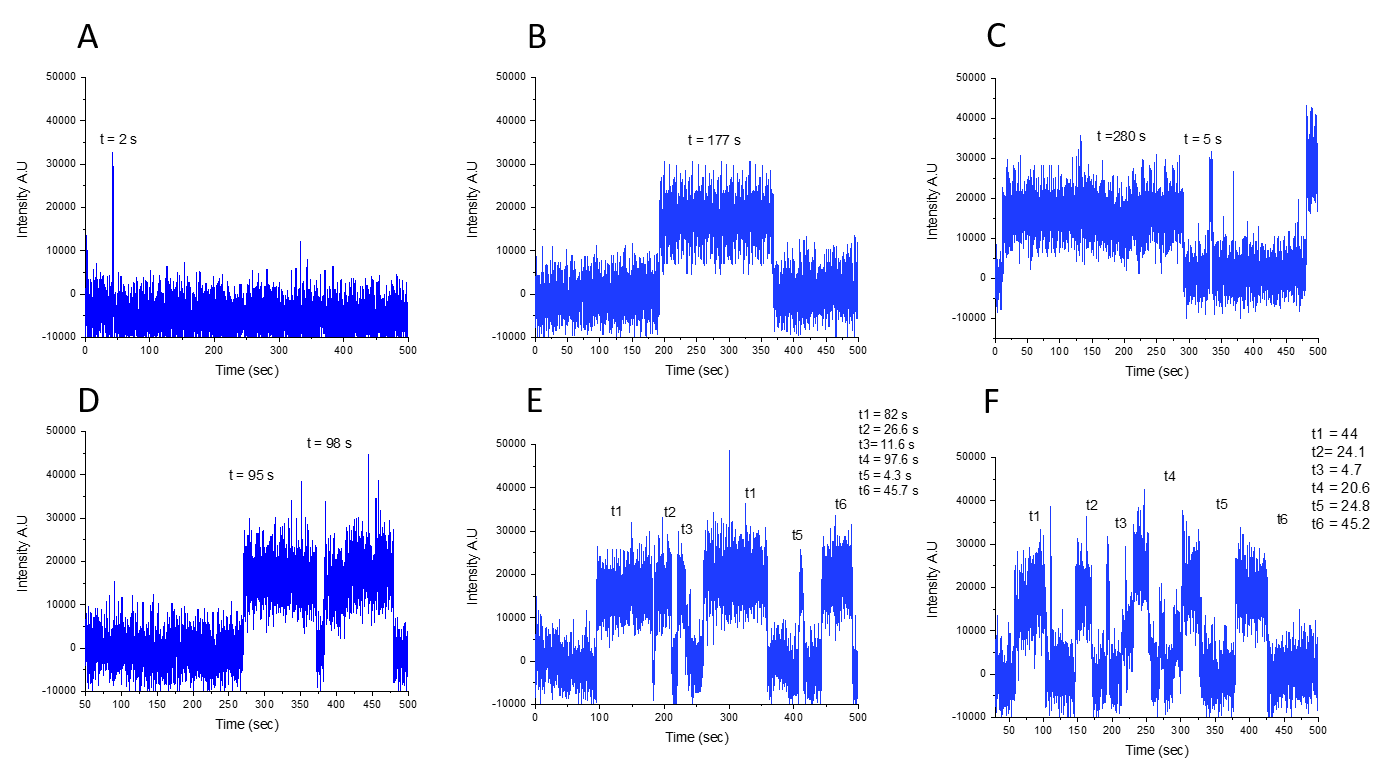
**Figure S7.** Example time trajectories of short and long lived SfiI-DNA complexes. The lifetime is obtained by the calculating the dwell time between the sudden rise and drop in the intensity from the base line. (A) Time trajectory with a dwell time of t = 2s; an example of the short lived SfiI-DNA complex. (B) and (C) The time trajectories with dwell times t = 177s, t = 280 s, and 5 s as examples of the long lived SfiI-DNA complexes. (D) Represents the time trajectory with a dwell time of t = 95s and t = 960 s, other examples of long lived SfiI-DNA complexes and the multiple association events on the same DNA duplex on the surface. (E) and (F) Represents the time trajectory with dwell time shown in the graph. This is an example of the SfiI-DNA complexes with the multiple association and dissociation of the complex to the same DNA duplex on the surface.

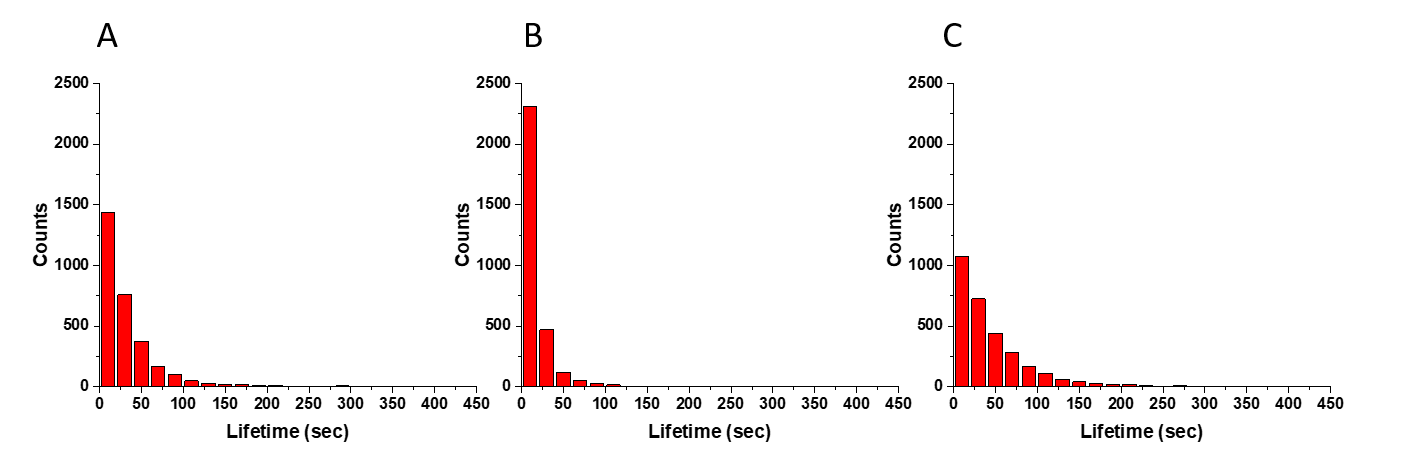

**Figure S8.** Frequency analysis of lifetimes of SfiI-DNA complexes with 23 bp DNA. (A) Represents the histogram of events for 23 bp synaptic complexes. (B) Represent the histogram of events for 23 bp non-specific complexes. (C) Represents the histogram of events for 23 bp pre-synaptic complexes with specific DNA tethered to the surface and nonspecific DNA labelled with Cy3.

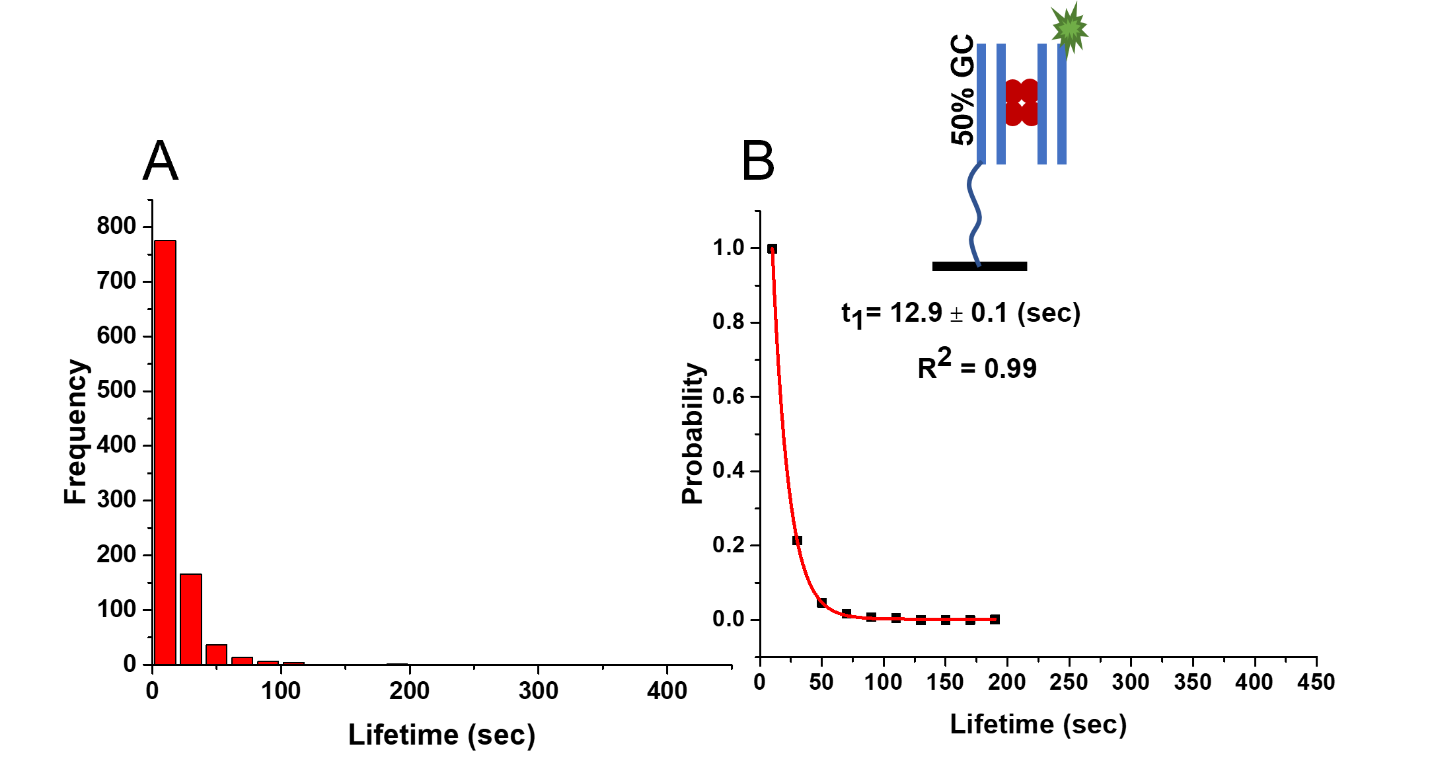

**Figure S9.** Lifetime of SfiI-DNA complexes with 23 bp with 50% GC content. (A) Histogram of SfiI-DNA complex lifetimes. (B) The normalized survival probability fit with a single exponential decay approximation. The characteristic lifetime of these complexes is 12.9 ± 0.1 (SD). The 23 bp DNA duplex is represented by two lines in blue. The Cy3 label is represented by a green star. The SfiI tetramer is represented with four red circles. The line drawn from the Duplex represents the tether attached to the surface to immobilize the duplex.

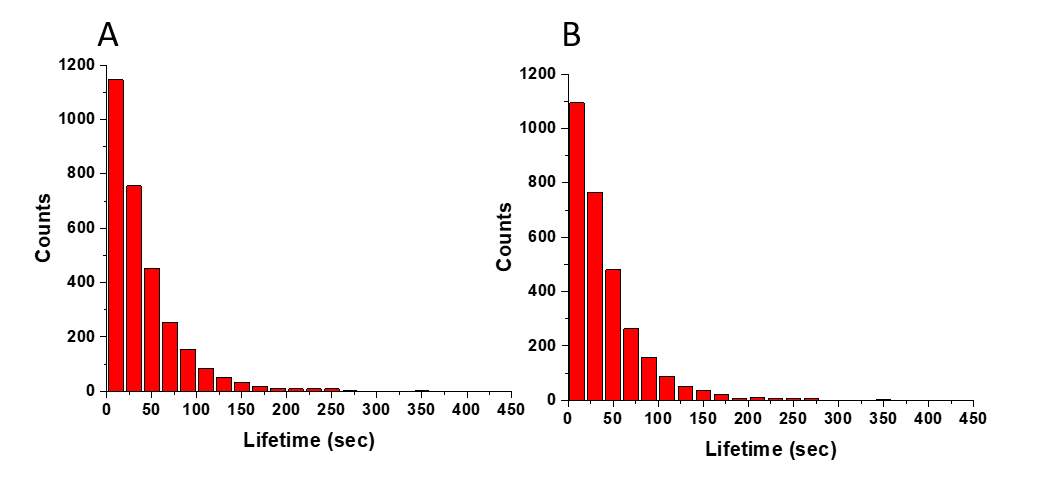

**Figure S10**. Frequency analysis of lifetimes of the interactions of the SfiI-DNA complexes formed using 23 bp DNA substrates. (A) Histogram for 23 bp pre-synaptic complexes, with non-specific DNA tethered to the surface. (B) Histogram for 23 bp pre-synaptic complexes with non-specific DNA with 50% GC content tethered to the surface.

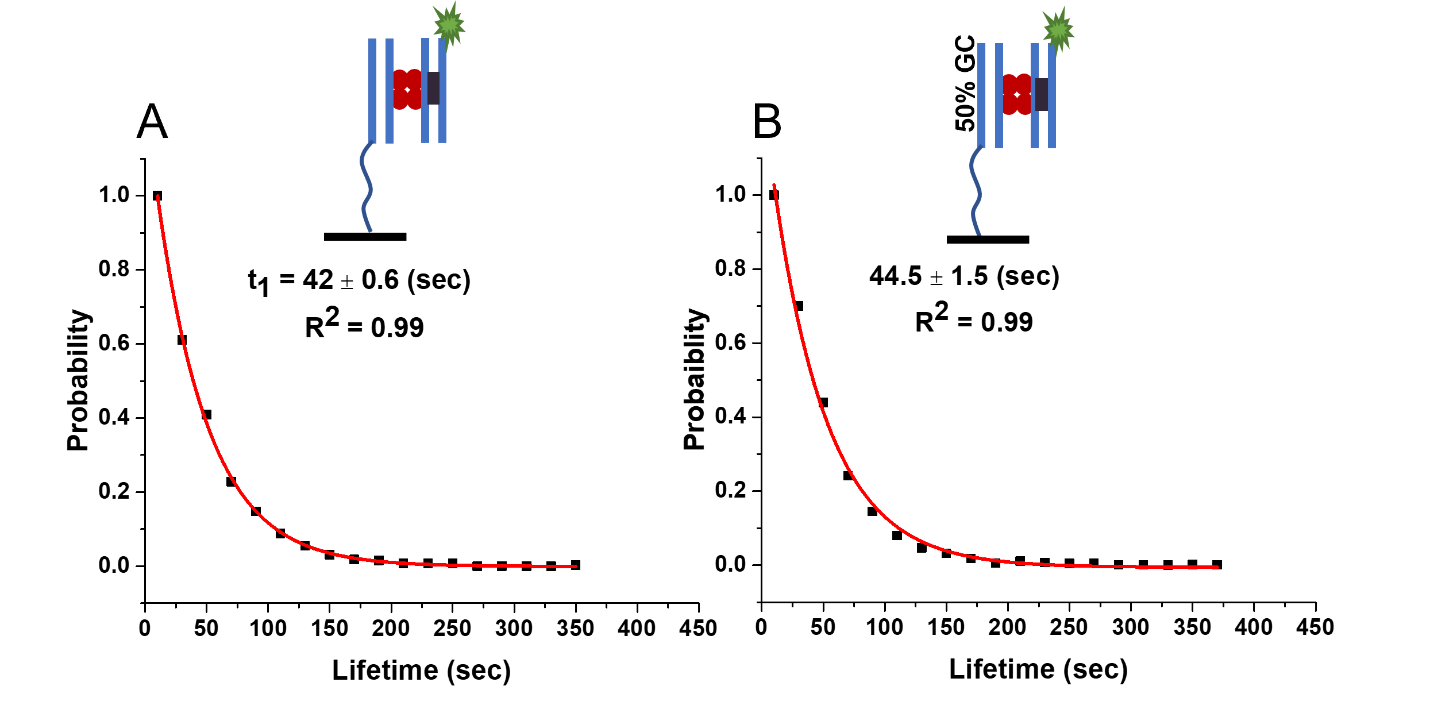

**Figure S11.** Normalized survival probability plots for Sfil-DNA complexes with 23 bp DNA. (A) Represents the lifetimes of the complexes formed between non-specific-specific (*ns*) duplexes (pre-synaptic complexes) with a characteristic lifetime of 42 ± 0.6 (sec ± SD), calculated using single exponential decay approximation. (B) Lifetime of the complexes formed between non-specific 23 bp duplex DNA with 50% GC content and 23bp specific (*ns)* duplexes (pre-synaptic complexes) with characteristic lifetime of 44.5 ± 1.5 (sec ± SD). with a single exponential decay fit. The 23 bp DNA duplex is represented by two lines in blue. The specific site for the SfiI is represented by a rectangle inside the blue lines. The Cy3 label is represented by a green star. The SfiI tetramer is represented with four red circles. The line drawn from the Duplex represents the tether attached to the surface to immobilize the duplex

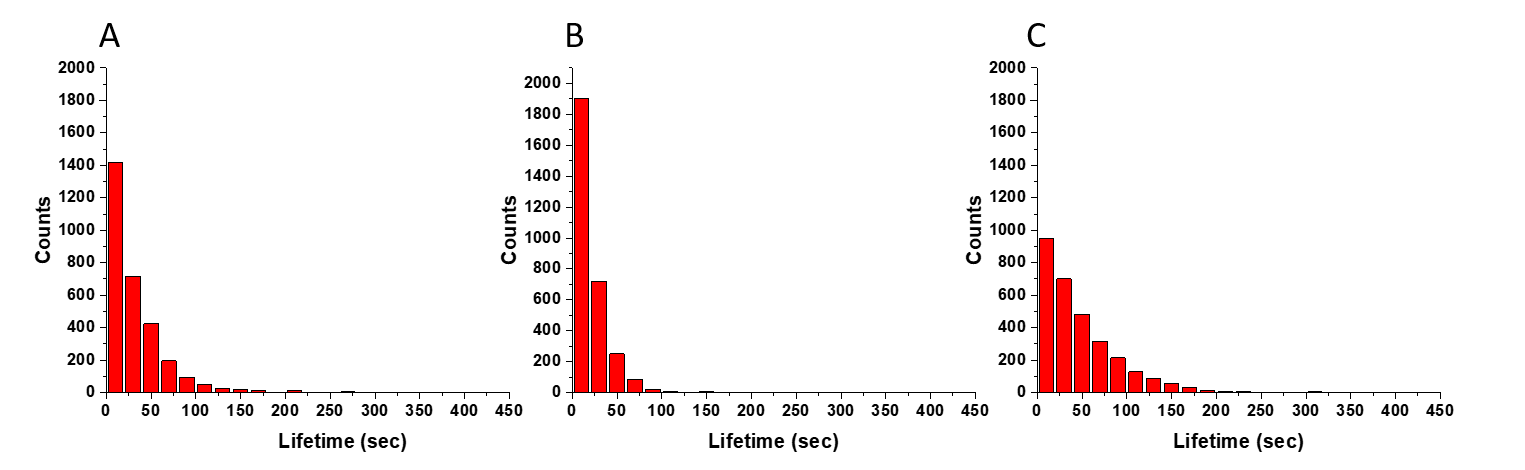

**Figure S12**. Frequency analysis of lifetimes of SfiI-DNA complexes with 33 bp DNA substrates. (A) Histogram of lifetimes for 33 bp synaptic complexes. (B) Histogram of lifetimes for 33 bp non-specific complexes. (C) Histogram of lifetime for 33 bp pre-synaptic complexes, with specific DNA tethered on the surface.

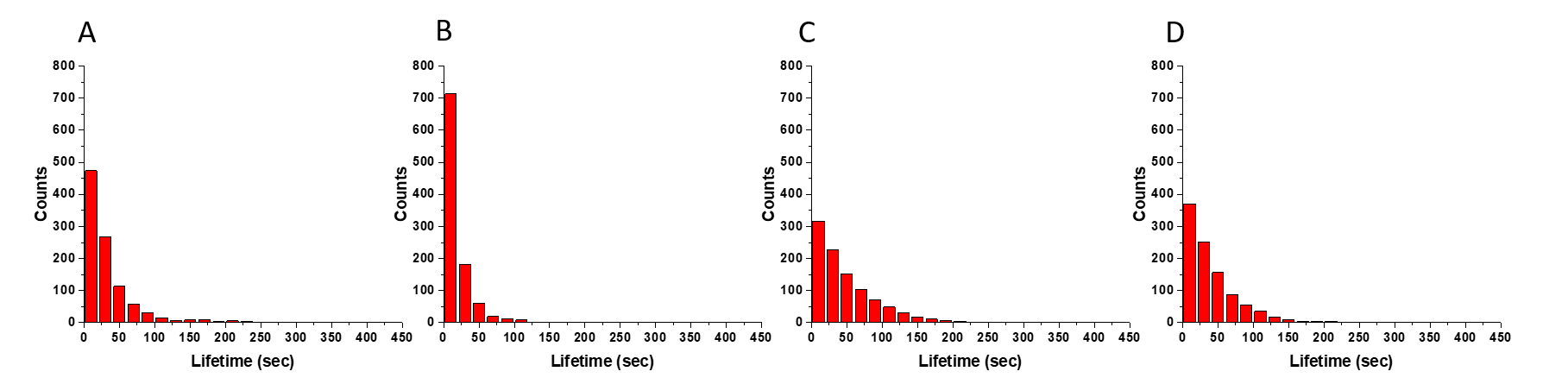

**Figure S13.** Frequency analysis of lifetimes of SfiI-DNA complexes formed using asymmetric DNA duplex lengths, with 23bp and 33 bp DNA. (A) Histogram for 23 bp specific (on surface) and 33 bp specific synaptic complexes. (B) Histogram for 23 bp (on surface) and 33 bp non-specific complexes. (C) Histogram for 23 and 33 bp (on surface) pre-synaptic complexes. (D) Histogram for 23 (on surface) and 33 bp pre-synaptic complexes.
